# ResistoXplorer: a web-based tool for visual, statistical and exploratory data analysis of resistome data

**DOI:** 10.1101/2020.08.14.250837

**Authors:** Achal Dhariwal, Roger Junges, Tsute Chen, Fernanda Cristina Petersen

## Abstract

The study of resistomes using whole metagenomic sequencing enables high throughput identification of resistance genes in complex microbial communities, such as the human microbiome. Over recent years, sophisticated and diverse pipelines have been established to facilitate raw data processing and annotation. Despite the progress, there are no easy-to-use tools for comprehensive visual, statistical, and functional analysis of resistome data. Thus, exploration of the resulting large complex datasets remains a key bottleneck requiring robust computational resources and technical expertise, which creates a significant hurdle for advancements in the field. Here, we introduce ResistoXplorer, a user-friendly tool that integrates recent advancements in statistics and visualization, coupled with extensive functional annotations and phenotype collection, to enable high-throughput analysis of common outputs generated from metagenomic resistome studies. ResistoXplorer contains three modules- the ‘Antimicrobial Resistance Gene Table’ module offers various options for composition profiling, functional profiling and comparative analysis of resistome data; the ‘Integration’ module supports integrative exploratory analysis of resistome and microbiome abundance profiles derived from metagenomic samples; finally, the ‘Antimicrobial Resistance Gene List’ module enables users to intuitively explore the associations between antimicrobial resistance genes and the microbial hosts using network visual analytics to gain biological insights. ResistoXplorer is publicly available at http://www.resistoxplorer.no.

## INTRODUCTION

Antimicrobial resistance (AMR) represents a major threat to global public health and the economy [1]. Consequently, examining the emergence and dissemination of AMR genetic determinants is one of the priorities in global research [2–4]. Until recently, genetic determinants were mostly understood in the context of specific pathogens. However, to fully understand how antimicrobial resistance genes (ARGs) emerge and disseminate a more holistic approach is required. In this respect, advancements in short-read based high-throughput DNA sequencing (HTS) technologies and computation methods have facilitated rapid identification and characterization of ARGs across all the microbial genomes present in a sample (metagenome) [5, 6]. They have been shown to provide unprecedented knowledge into the large reservoir of ARGs and contributed to elucidate the ARG composition and the spread of AMR between human, animal, and environmental microbial communities [7–12]. Currently, resistome profiles describing ARGs in complex and diverse microbial metagenomes are primarily generated using whole metagenome shotgun sequencing in which the total DNA extracted from a microbial community is sequenced. The resulting DNA fragments can be analyzed using read or assembly-based approaches to characterize their resistome composition [5, 6]. These derived sequencing datasets are both large and complex, causing considerable ‘big data’ challenges in downstream data analysis.

The main computational effort in resistome analysis of metagenomic datasets so far has focused on processing, classification, assembly, and annotation of sequenced reads. This has led to the development of a number of excellent bioinformatic tools and databases for detecting and quantifying ARGs in metagenomes [5, 6, 13, 14]. However, there is still no clear consensus with regards to standard analysis pipelines and workflows for high-throughput analysis of AMR metagenomic resistome data [14, 15]. Nonetheless, the outputs from most of these pipelines can be summarized as a data table consisting of feature (ARGs) abundance information across samples, *i.e*. resistome profiles, along with their functional annotations and sample metadata. For most researchers, the fundamental challenge in data analysis can often be centered on how to understand and interpret the information in the abundance tables especially within the context of different experimental factors and annotations.

Downstream analysis of resistome data can be separated into four main categories: (i) composition profiling-to visualize and characterize the resistome based on approaches developed in community ecology such as alpha diversity, rarefaction curves or ordination analysis; (ii) functional profiling- to analyze resistome profiles at different functional categories (*e.g*. Drug class, Mechanism), thus gaining better insights on their collective functional capabilities; (iii) comparative analysis- to identify features having a significant differential abundance between studied conditions, and (iv) integrative analysis- to integrate the resistome and taxonomic data to understand the complex interplay and potential associations between microbial ecology and AMR. The computational methods and approaches to perform such analysis are fairly diverse and require deep understanding and programming skills, representing significant barriers for their broader and exploratory applications [16]. The first category of analysis can be more straightforward to perform, but the last three are challenging.

Fundamental challenges within the different categories relate to the fact that metagenomic data is often characterized by differences in library sizes, sparsity, over-dispersion, and compositionality [17, 18]. Hence, it is critical to normalize the data to achieve comparable and meaningful results [18–20]. To deal with uneven library sizes, researchers often employ two common normalization approaches prior to analysis: subsampling the reads in each sample to the same number (rarefying) or rescaling the total number of reads in each sample to a uniform sum (using proportions). The former may entail loss of valuable information, while the latter could lead to issues related to data compositionality [21]. To overcome such challenges, sophisticated scaling methods based on log-ratio transformations have been proposed [22, 23]. To identify differentially abundant genes, the development of statistical models that account for features of metagenomic data or the use of methods to transform data to have distributions that fit standard test assumptions is generally recommended [19]. For instance, the metagenomeSeq algorithm incorporates cumulative sum scaling normalization and a zero-inflated Gaussian (ZIG) mixture model to reduce false positives and improve statistical power for differential abundance analysis [24, 25]. It has also been demonstrated that algorithms developed for RNA-seq data such as edgeR and DESeq2, along with their respective normalization methods, outperform other approaches used for metagenomic abundance data [20, 25, 26]. These standard strategies are widely employed, but do not explicitly account for the compositional nature of whole metagenomic sequencing data [27, 28]. To address this issue, promising Compositional Data Analysis (CoDA) approaches have been proposed [29, 30].

Nonetheless, there is no one statistical method suitable for all types of metagenomic datasets [20, 26]. The best results are achieved from a trade-off between data characteristics (sample or group size, sequencing depth, effect sizes, genes abundances, etc.) and the normalization method, incorporated with the coupled exploratory and comparative analysis [31]. Therefore, various statistical and normalization methods are required for different metagenomic datasets, analyses and research questions addressed [6, 19, 25, 26, 31]. However, most of the approaches have been implemented as R packages. Although flexible, learning R in order to use these methods can be challenging for most clinicians and researchers.

In the second category, functional profiling, the characterized resistome abundance profiles are analyzed by mapping ARGs either to their respective class of drugs to which they confer resistance (Class-level) or to their underlying molecular mechanism of resistance (Mechanism-level). Analyzing resistomes at such high level categories enables researchers to gain more biological, actionable, and functional insights together with a better understanding of their data. However, these functional levels and categories, along with their classification scheme, vary considerably between AMR reference databases [15]. Additionally, depending upon the database used for annotation, users need to manually collect and curate such information and then generate separate abundance tables for each functional level. Hence, collecting appropriate functional annotation information for hundreds of ARGs in resistomes for functional profiling and further downstream analysis can be confusing, arduous, time-consuming, and error-prone. On the contrary, some of these databases may also provide information regarding the microbial hosts that harbor or carry these reference ARGs. Information about such relationships can be complex as one microbe can carry multiple ARGs and single ARGs can in turn be present across multiple microbes. To explore such intricate ‘multiple-to-multiple’ relations, one option is to use a network-based visualization method. However, such visual exploration support is not present in current resistome analysis tools.

To address these gaps as well as to meet recent advances in resistome data analysis, we have developed ResistoXplorer, a user-friendly, web-based, visual analytics tool to assist clinicians, bench researchers, and interdisciplinary groups working in the AMR field to perform exploratory data analysis on abundance profiles and resistome signatures generated from AMR metagenomics studies. The key features of ResistoXplorer include:

- Support of a wide array of common as well as advanced methods for composition profiling, visualization and exploratory data analysis;
- Comprehensive support for various data normalization methods coupled with standard as well as more recent statistical and machine learning algorithms;
- Support of a variety of methods for performing vertical data integrative analysis on paired datasets (*i.e*. taxonomic and resistome abundance profiles)
- Comprehensive support for ARG functional annotations along with their microbe and phenotype associations based on data collected from >10 reference databases;
- A powerful and fully featured network visualization for intuitive exploration of ARG-microbe associations, including functional annotation enrichment analysis support.

Collectively, these features consist of a comprehensive tool suite for exploratory downstream analysis of data generated from AMR metagenomics studies. ResistoXplorer is freely available at http://www.resistoxplorer.no.

## MATERIAL & METHODS

### Design & Implementation

ResistoXplorer is implemented based on Java, R and JavaScript programing languages. The framework is developed based on the Java Server Faces technology using the PrimeFaces (https://www.primefaces.org/) and BootsFaces (https://www.bootsfaces.net) component library. The network visualization uses the sigma.js (http://sigmajs.org/) JavaScript library. Additionally, D3.js (https://d3js.org/) and CanvasXpress (https://canvasxpress.org/) JavaScript libraries are utilized for other interactive visualization. All the R packages for performing back-end analysis and visualization are mentioned in the ‘About’ section of the tool. At the start of the analysis, a temporary account is created with an associated home folder to store the uploaded data and analysis results. All the analysis results will be returned in real-time. Upon completing their analysis session, users should download all their results. The system is deployed on a dedicated server with 4 physical CPU cores (Intel Core i5 3.4GHz), 8GB RAM and Ubuntu 18.04 LTS was used as the operation system. ResistoXplorer has been tested with major modern browsers such as Google Chrome, Mozilla Firefox, Safari, and Microsoft Internet Explorer.

### Program description and methods

ResistoXplorer consists of three main analysis modules. The first is the **ARG List** module that is designed to explore the functional and microbial hosts associations for a given list of antimicrobial resistance genes (ARGs) of interest. The second is the **ARG Table** module, which contains functions for analyzing resistome abundance profiles generated from AMR metagenomics studies. Lastly, the **Integration** module enables users to perform integrative analysis on the paired taxonomic and resistome abundance profiles to further explore potential associations coupled with novel biological insights and hypotheses. Figure 1 represents the overall design and workflow of ResistoXplorer. We recommend users to try out our example datasets to get familiar with the basic steps and key features of the tool before proceeding with analysis of their own data. ResistoXplorer also contains manuals and a comprehensive list of frequently asked questions (FAQs) to assist users to easily navigate through different analysis tasks.

**Figure 1:**
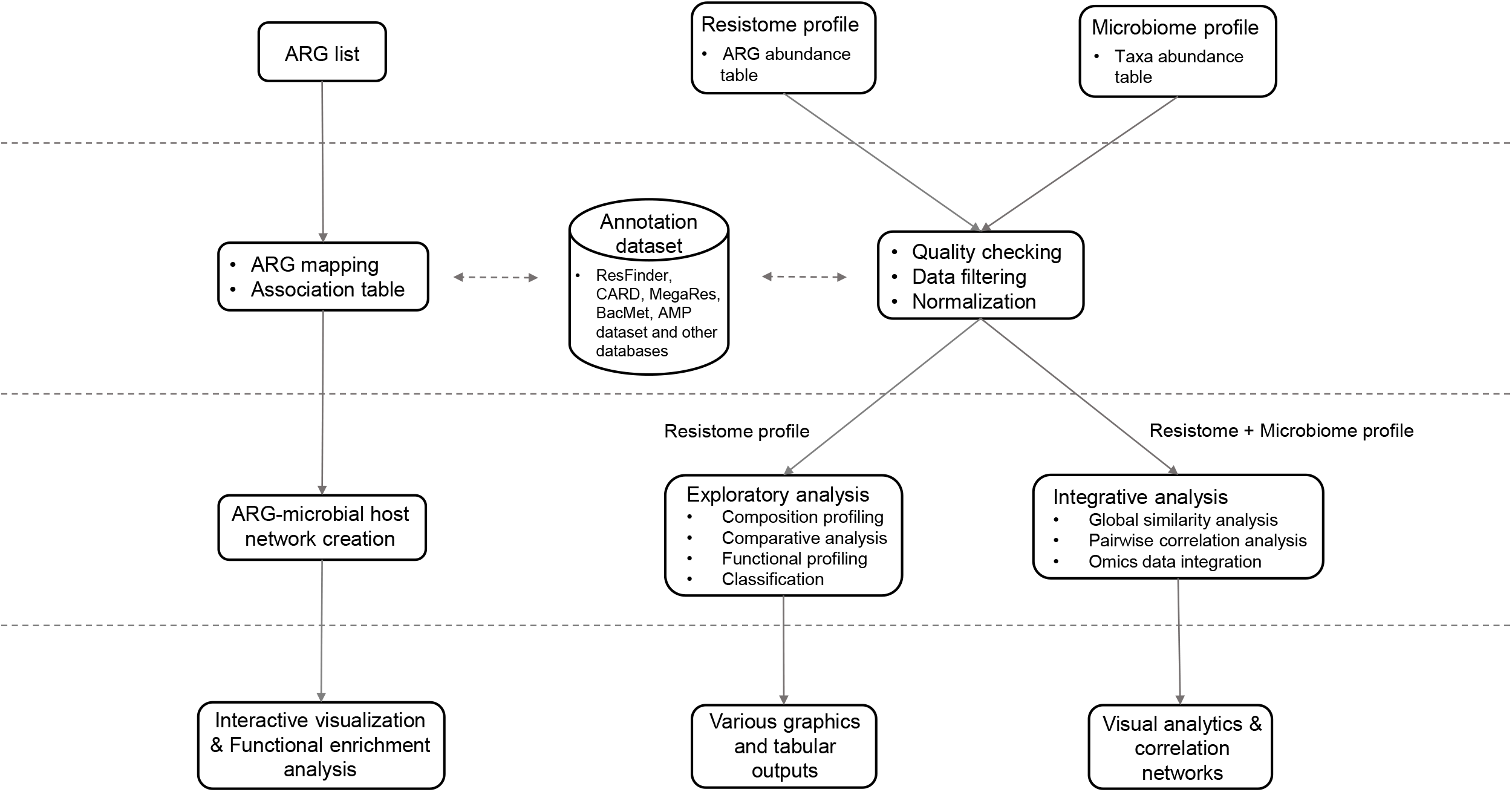
ResistoXplorer flow chart. ResistoXplorer accepts resistance gene list and ARG/taxa abundance tables as input data. Three successive steps are performed - data processing, data analysis, and result exploration. The accompanying web interface proffers a varied suite of options, and generates several tables and graphics to enable users to intuitively go over the data analysis tasks.

## RESULTS & DISCUSSION

### Data upload and processing

#### Overview of data inputs

The three analysis modules (ARG List, ARG Table and Integration) are represented as three interactive buttons in ResistoXplorer. Users must choose an analysis path based on their input. The input can be uploaded in two different ways – by entering a list of antimicrobial resistance genes (ARGs) or by uploading an ARG abundance table along with a sample metadata file containing group information. In the latter case, the files can be uploaded as a tab-delimited text (.txt) or in comma-separated values (.csv). Further, users must also provide the annotation information of ARGs either by uploading a file (.txt or .csv) or by just selecting the same database used for their annotation during upstream analysis. Additionally, the Integration module requires users to also upload a taxa abundance table in the same formats. The taxonomic and functional annotation files are optional in case of integrative analysis. Users can go to the corresponding ‘Manuals’ and ‘Data Format’ section, or download the example datasets for more details.

#### ARG-functional annotations collection

The annotation information and the classification scheme for reference ARGs (or sequences) are collected from 9 widely used generalized AMR databases: CARD [32], ResFinder [33], MEGARes [15, 34], AMRFinder [35], SARG [36], DeepARG-DB [37], ARGminer [38], ARDB [39] and ARG-ANNOT [40]. Further, this annotation information is organized into tables containing the reference ARGs in rows and their functional annotation levels across columns for each of the databases. The headers (names) for reference ARGs are annotated as in the chosen database. Some of the annotation levels having multiple functional category assignments for ARGs were removed from the tables to avoid false counts inflation during downstream analysis. Additional manual curation for functional category naming and data redundancies were performed in some of the databases. Further, we also structured functional annotation information from the BacMet [41] database and antimicrobial peptide (AMP) resistance gene dataset [42] to enable users to perform functional profiling and downstream analysis of antibacterial biocides/metals and AMP resistance genes abundance profiles. It should be noted that ResistoXplorer does not perform any functional annotation of sequencing data. The tables are stored in RDS file format for less storage space and fast retrieval of data. Users can use these tables to analyze their resistome profile directly at different functional levels rather than manually collecting the annotation information from their corresponding database. The ‘Data Format’ and ‘About’ pages provide a detailed description on the format, structure of annotation table and database statistics, together with the links to allow users to download the annotation structure (‘Downloads’ section) available for the individual database.

#### Data filtration and normalization

By default, features with zero read count across all the samples or only present in one sample with extremely low count (*e.g*., 2) are removed from downstream analysis based upon some statistical and biological approximations. Also, features present in only a small percentage of samples (*e.g*., 20%) with very few counts (*e.g*., 2) cannot be discriminated from sequencing errors or low-level contamination, and also considered difficult to interpret their significance with respect to the whole community. By default, such low-quality features are filtered based on sample prevalence and their abundance levels to improve the comparative analysis. The default values are those used by the other tools and mostly found in the literature [43, 44]. Users can also choose to remove low abundant features by setting a minimum count cutoff based on their mean or median value. Conversely, some features remain constant in their abundances throughout the experimental conditions or across all the samples. These features are implausible to be informative in the comparative analysis. Users can exclude such low variant features based on their inter-quantile ranges, standard deviations or coefficient of variations [43]. Removing those uninformative features can increase the statistical power by reducing multiple testing issues during differential analysis [45]. The filtered data is used for most of the downstream analysis except alpha diversity and rarefaction analysis. In case of integrative analysis, users can also choose to apply different data filtration criteria for both taxonomic and resistome abundance data.

After data filtering, users must perform data normalization to remove the systematic variability between samples. Currently, ResistoXplorer offers three categories of data normalization-rarefying, scaling and transformation, based on various widely used methods for metagenomic abundance data [25]. In addition, ResistoXplorer supports other normalization methods like centered log-ratio (clr) and additive log-ratio (alr) transformation to facilitate compositional data analysis. The choice of method is dependent upon the type of analyses to be performed [20, 31]. The normalized data is used for exploratory data analysis including ordination, clustering and integrative analysis. Users can explore different approaches and visually investigate the clustering patterns (*i.e*., ordination plots, dendrogram and heatmap) to determine the effects of different normalization methods with regard to the experimental factor of interest. The total sum scaling (TSS) normalization is recommended for such type of analysis and has been set as the default option in ResistoXplorer [20, 46, 47]. Also, comparative analysis using different approaches is performed on normalized data. However, each of these approaches will use its own specific normalization procedure due to the lack of benchmark study evaluating which normalization methods should be combined with the different statistical approaches to achieve best performance for identifying differentially abundant genes [26]. For example, the relative log expression (RLE) normalization is used for DESeq2, and the centered log-ratio transformation is applied for ALDEx2. In the integrative analysis module, taxonomic and resistome datasets are normalized using the same approach.

### Composition profiling

#### Visual exploration

ResistoXplorer allows users to visually explore the resistome based on various intuitive visualization approaches used for metagenomic data. For instance, users can visualize resistome abundance data while simultaneously showing the functional hierarchical relationships and connectivity between features using an interactive sankey diagram, zoomable sunburst or treemap graphics (Figure 2C). In case of treemap, users can click a particular rectangular block of interest to further inspect its compositions at a lower functional level. The abundances can be represented as either absolute counts or relative proportions. However, such visualizations are more suitable for resistome profiles having an acyclic and hierarchical functional annotation structure. Abundance profiles can also be viewed at different functional levels using other common visualizations such as stacked area or stacked bar plot (Figure 2A). The plot is organized by experimental factors to help visualize the differences in resistome composition across different conditions.

**Figure 2:**
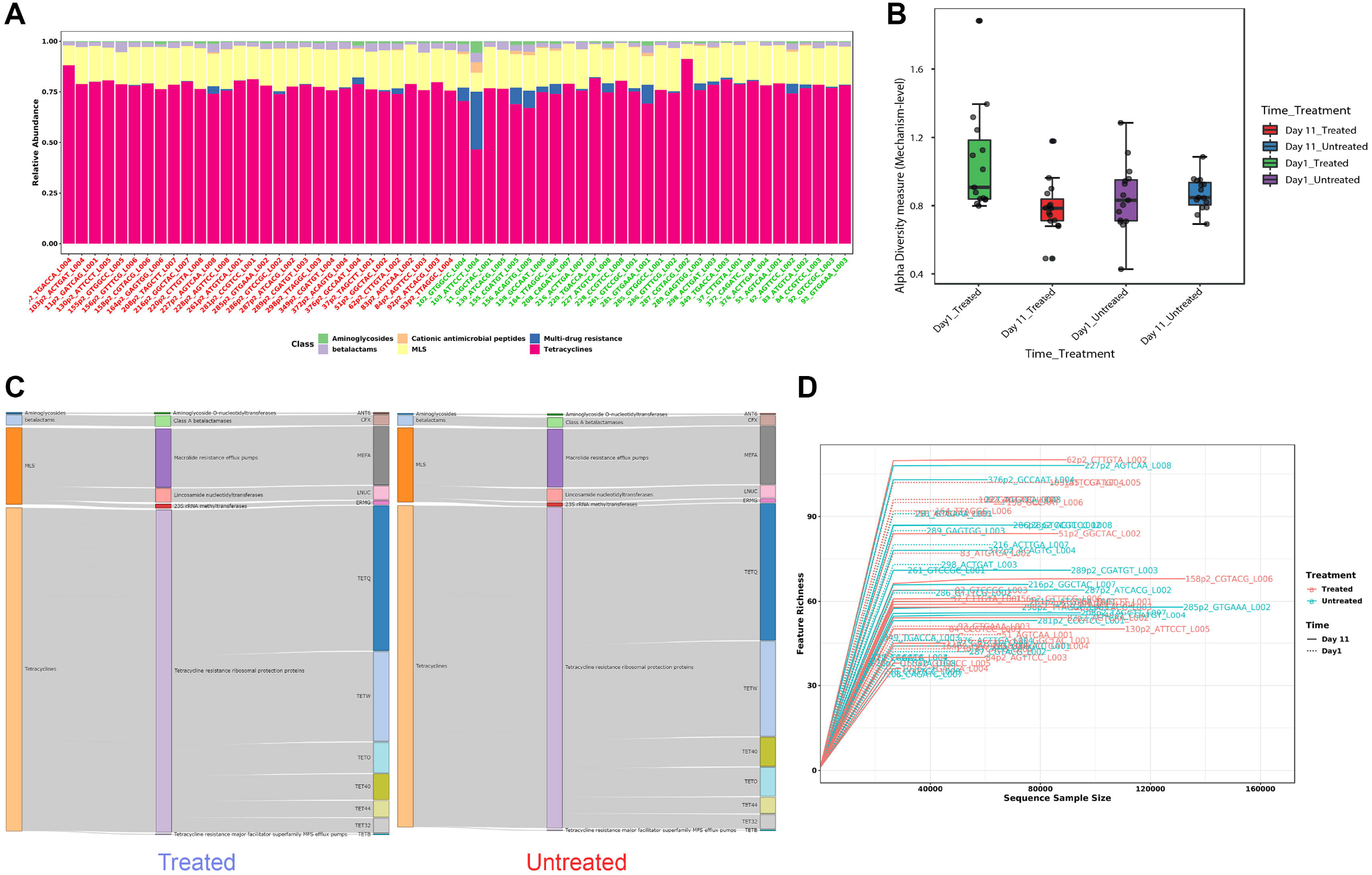
Example outputs from composition profiling panel of ARG Table module in ResistoXplorer. (A) A stacked bar chart showing class level resistance abundance profiles across samples. (B) A box plot summary of the Shannon diversity index at mechanism level in different treatment groups across sampling time points. (C) A Sankey diagram showing the resistome abundance profile of treated (left) and untreated (right) cattle group at hierarchical functional levels including class, mechanism and group. (D) A rarefaction curve showing the number of unique ARGs identified in each sample as a function of sequence sample size.

#### Diversity profiling

The resistome diversity profiling is implemented mainly based on the R vegan and phyloseq packages [48, 49]. Currently, users can perform alpha diversity (within-sample) analysis using eight common richness and/or evenness-based diversity measures. Since the Chao1 measure performed well and is recommended for estimating ARG diversity [50], it has been set as a default one in ResistoXplorer. The results are represented in the form of a dot plot for individual samples and box plots for each sample group (Figure 2B). The corresponding statistical significance is calculated automatically using either parametric or non-parametric tests. The analysis can be performed at different functional levels based on the available annotations. Additionally, the reliability of estimated diversity in samples can be assessed through rarefaction curves in which the number of unique features (ARGs) identified is plotted against the sequence sample size (Figure 2D).

#### Ordination analysis

The ordination analysis function allows users to explore and visualize the similarities or dissimilarities between samples based on their composition at different functional levels. The dissimilarity can be calculated using five non-phylogenetic-based quantitative or qualitative distance measures. Since the different types of distance measures have specific niches and can affect the outcomes and the interpretation of the analysis, it is recommended by several authors to apply different measures to better understand the factors underlying composition differences [6, 51, 52]. Currently, three widely accepted ordination methods are supported, including principal coordinate analysis (PCoA), non-metric multidimensional scaling (NMDS) and principal component analysis (PCA). In particular, users can follow a CoDA ordination approach by performing PCA on the centered log-ratio transformed data. The corresponding statistical significance is calculated using one of the three common multivariate statistical testing methods with generally the more powerful method i.e., Permutational multivariate analysis of variance (PERMANOVA) set as the default option [53]. By default, ordination analysis is performed using PCoA on a most widely used Bray–Curtis dissimilarity metric and assessed using PERMANOVA. The results are represented as both 2D and 3D sample plots. The samples visualized in these ordination plots are colored based on metadata or their alpha diversity measures to help users identify any underlying patterns in the datasets (Figure 3B).

**Figure 3:**
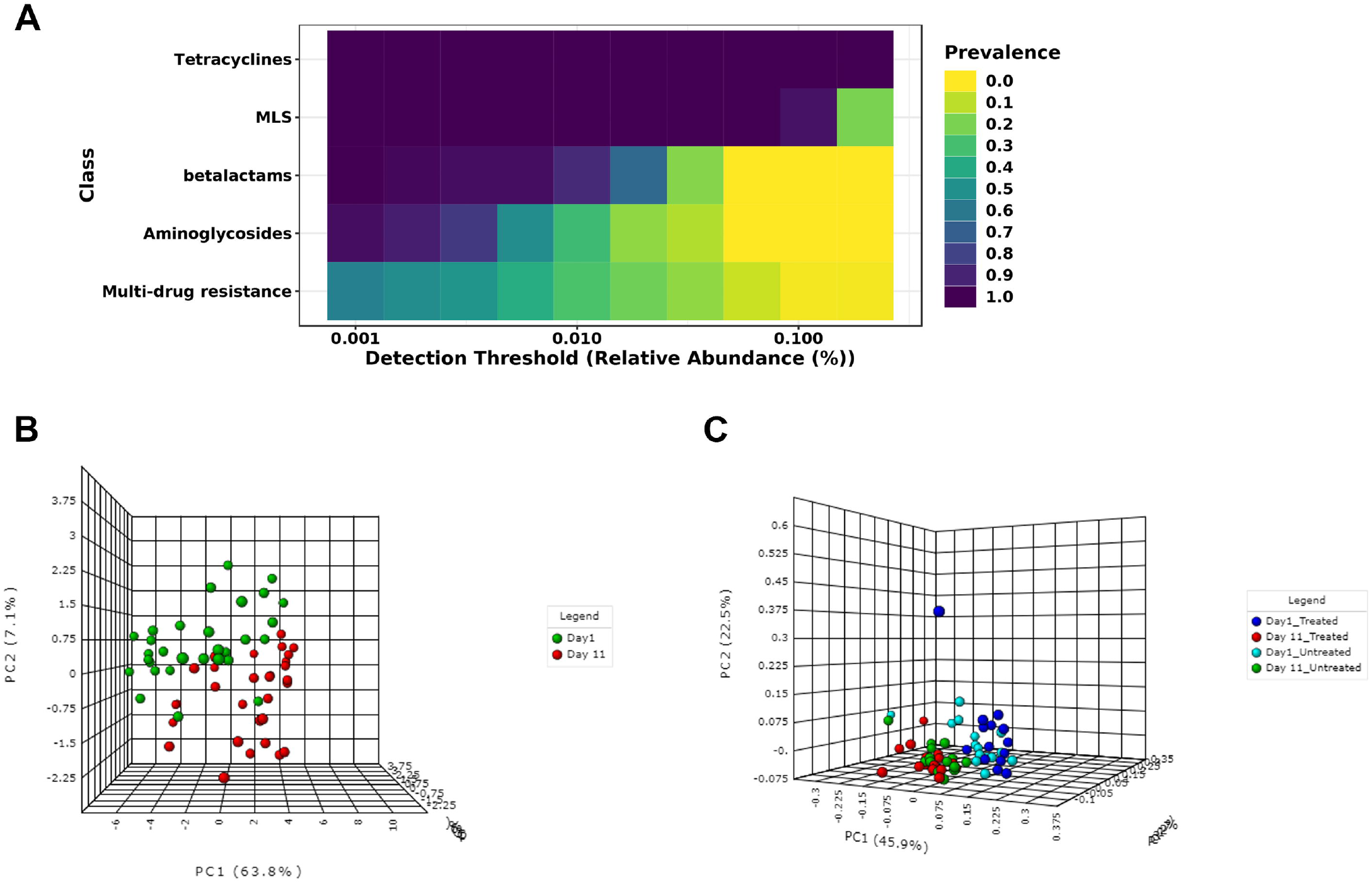
Illustration of core resistome and ordination analysis results in ResistoXplorer. (A) A heatmap showing the core resistome of cattle analyzed at class level. (B) A 3D PCA plot with sample colors based on time points. (C) A 3D PCoA plot with sample colors with regards to different treatment groups and time points.

### Comparative analysis

#### Differential abundance testing

This section enables users to perform statistical testing to identify features that are significantly different in abundance across sample groups. ResistoXplorer supports standard tests, such as DESeq2 [22], edgeR [23], metagenomeSeq [24], LEfSe [54], as well as more recent CoDA univariate analysis approaches such as ALDEx2 [55] and ANCOM [56]. DESeq2 and edgeR are broadly used and generally considered as robust and powerful parametric statistical approaches for datasets with small group and equally distributed library sizes [18, 20, 25, 26]. They fit a generalized linear model and assume that read counts follow a negative binomial distribution to account for the features of count data. In contrast, the metagenomeSeq with its recommended CSS normalization has substantially higher performance with larger group sizes [26, 48]. LEfSe uses the standard non-parametric tests for statistical significance coupled with linear discriminant analysis to assess the effect size of those differentially abundant features. The CoDA methods perform statistical testing on the log ratios of features rather than their actual count abundances to deal with the compositional nature of sequencing data. ALDEx2, for instance, performs parametric or nonparametric statistical tests on log-ratio values from a modeled probability distribution of the data and returns the expected values of statistical tests along with effect size estimates. ANCOM tests the log-ratio abundance of all pairs of features for differences in means using nonparametric statistical tests. By default, the Benjamini–Hochberg correction is used for all approaches to control the false discovery rate (FDR) across significant genes. The differential analysis can also be performed at different functional levels.

The results from the differential analysis are displayed as a table. By default, the result table will show a maximum of 500 top features ordered according to their adjusted P-values. The significant features (if present) are automatically highlighted in orange color. Further, users can also see a boxplot summary for any feature of interest by just clicking the ‘Details’ icon. Since different statistical approaches may generate divergent P-values, it is often recommended to compare and visualize results using more than one statistical approach, as to increase confidence in the interpretation of results.

#### Machine learning-based classification

Prediction of microbiome signatures using machine learning algorithms has been gaining more recognition and shown to perform well in recent resistome data analyses and classifications [7, 57, 58]. ResistoXplorer provides two such powerful supervised classification methods - Random Forest [59] and Support Vector Machine (SVM). Both can be applied to resistome data for identification of potential biomarkers. In particular, the Random Forest algorithm uses an ensemble of classification trees (forest), with final class prediction based on the majority vote of the ensemble. As the forest is constructed, it can provide an unbiased estimate of prediction errors by aggregating cross-validation results using bootstrapped samples. Random forest also measures the importance of each feature based on the increase of the prediction errors when it is randomly shuffled. Alternatively, the SVM algorithm uses a training set of samples separated into classes to identify a hyperplane in higher dimensional feature space that generates the largest minimum distance (margin) between the samples that belongs to different classes [60]. ResistoXplorer’s SVM analysis is performed using recursive feature selection and sample classification via linear kernel [61]. The features used by the best model are considered to be important and ranked based on their frequencies of being selected in the model. Figure 4D shows the classification performance of SVM with regards to decreasing number of features (variables).

**Figure 4:**
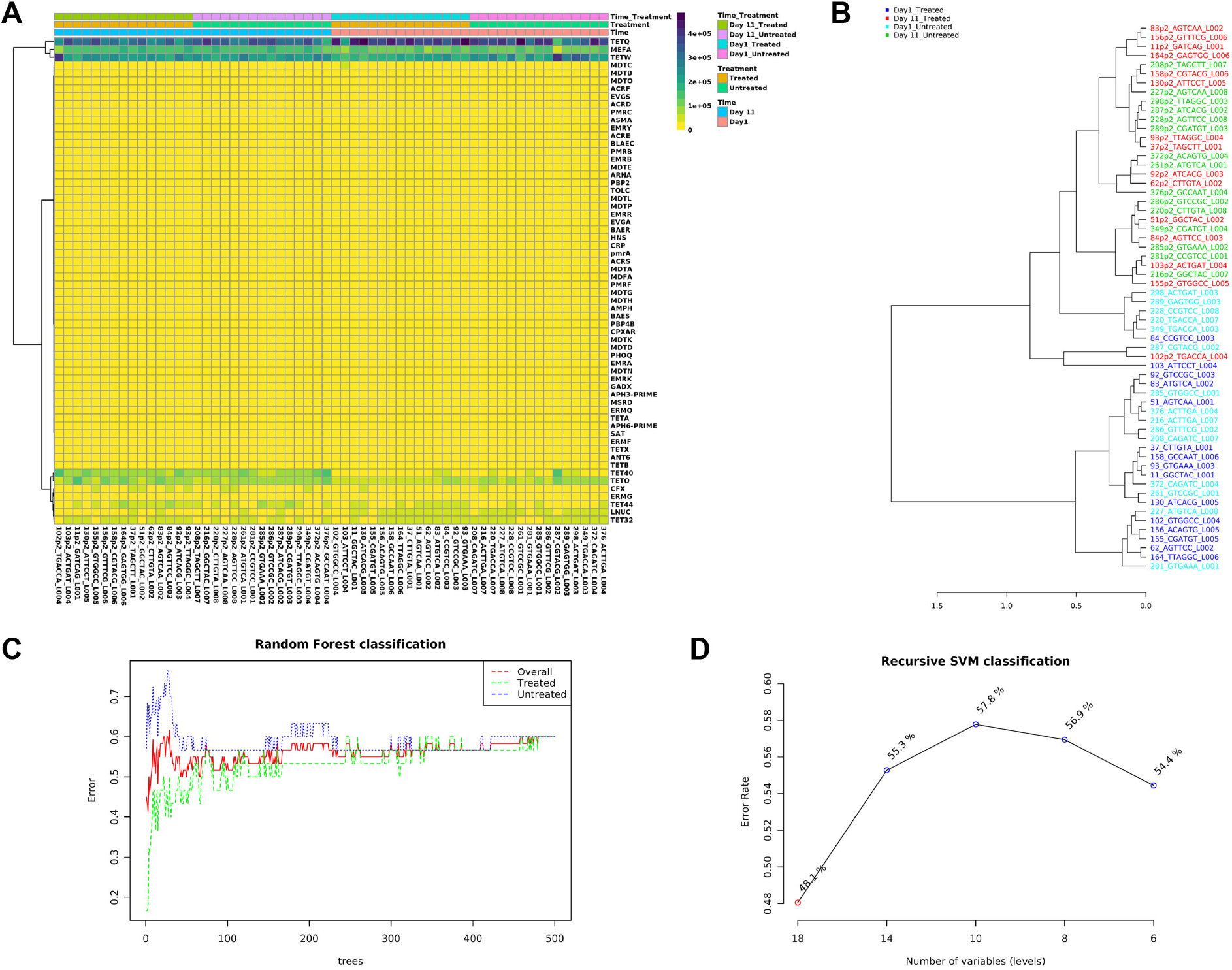
Example outputs from clustering analysis and machine learning-based classifications in ARG Table module of ResistoXplorer. (A) A clustered heatmap showing the variation of resistome abundance at group level in samples organized based on time point. (B) A dendrogram showing the clustering of samples with colors based on treatment and time point. (C) and (D) A graphical summary of the classification performance on different treatment groups using the Random Forests and SVM algorithm, respectively.

#### Other features

There are several other valuable functions implemented in ResistoXplorer for exploratory analysis of resistome data. Users can perform core resistome analysis to detect core sets of features present in samples or sample groups based on their abundance and prevalence level (Figure 3A). ResistoXplorer also supports commonly used correlation analysis and hierarchical clustering. The results from hierarchical clustering can be visualized using heatmaps (Figure 4A) and dendrograms (Figure 4B). For publication purposes, all visualizations can be downloaded as either Scalable Vector Graphics (SVG) or Portable Document Format (PDF) files.

### Integrative analysis

The main goal of this module is to explore and unveil potential hidden correlations between the microbiome and resistome using a variety of integrative data analysis approaches. Such analyses have been increasingly used to explore the associations between the bacteria and ARGs in different environments [11, 62, 63]. Currently, ResistoXplorer offers several advanced and commonly used univariate and multivariate statistical methods for data integration and correlation analysis. All these analyses are performed on filtered and normalized datasets.

#### Global Similarity analysis

This section allows users to determine the overall similarity between the microbiome and resistome dataset using two multivariate correlation-based approaches - Procrustes analysis (PA) and Coinertia analysis (CIA). The datasets used for such analysis can be normalized using scaling and/or transformation approaches to account for uneven library sizes and compositional data. The analysis can be performed at various functional and taxonomic levels based on available annotations. These functions currently support five common distance measures. The ordinations from the distance matrices can be calculated using either PCoA, PCA or NMDS. By default, both these analyses are being assessed employing widely used PCoA on a Bray–Curtis distance metric. In case of PA, the microbiome ordination is scaled and rotated onto the resistome ordination to minimize the sum of squared differences between the two ordinations. For the CIA, the microbiome and resistome ordination are constrained so that the squared covariances between them are maximized to measures the congruence between datasets. The results are represented using both 2D and 3D ordination plots, where samples are colored and shaped based on the datasets (Figure 5A). Users can also color the samples based on different experimental factors to identify some patterns or gain biological insights. The corresponding similarity coefficient and P-value from both these analyses are estimated automatically to assess the strength and significance of the association between the two datasets. The similarity coefficient ranges from 0 and 1, with 0 suggesting total similarity and 1 total dissimilarity between the two datasets. Users can perform both Procrustes and Coinertia analysis to gain more confidence by evaluating the congruency of the results.

**Figure 5:**
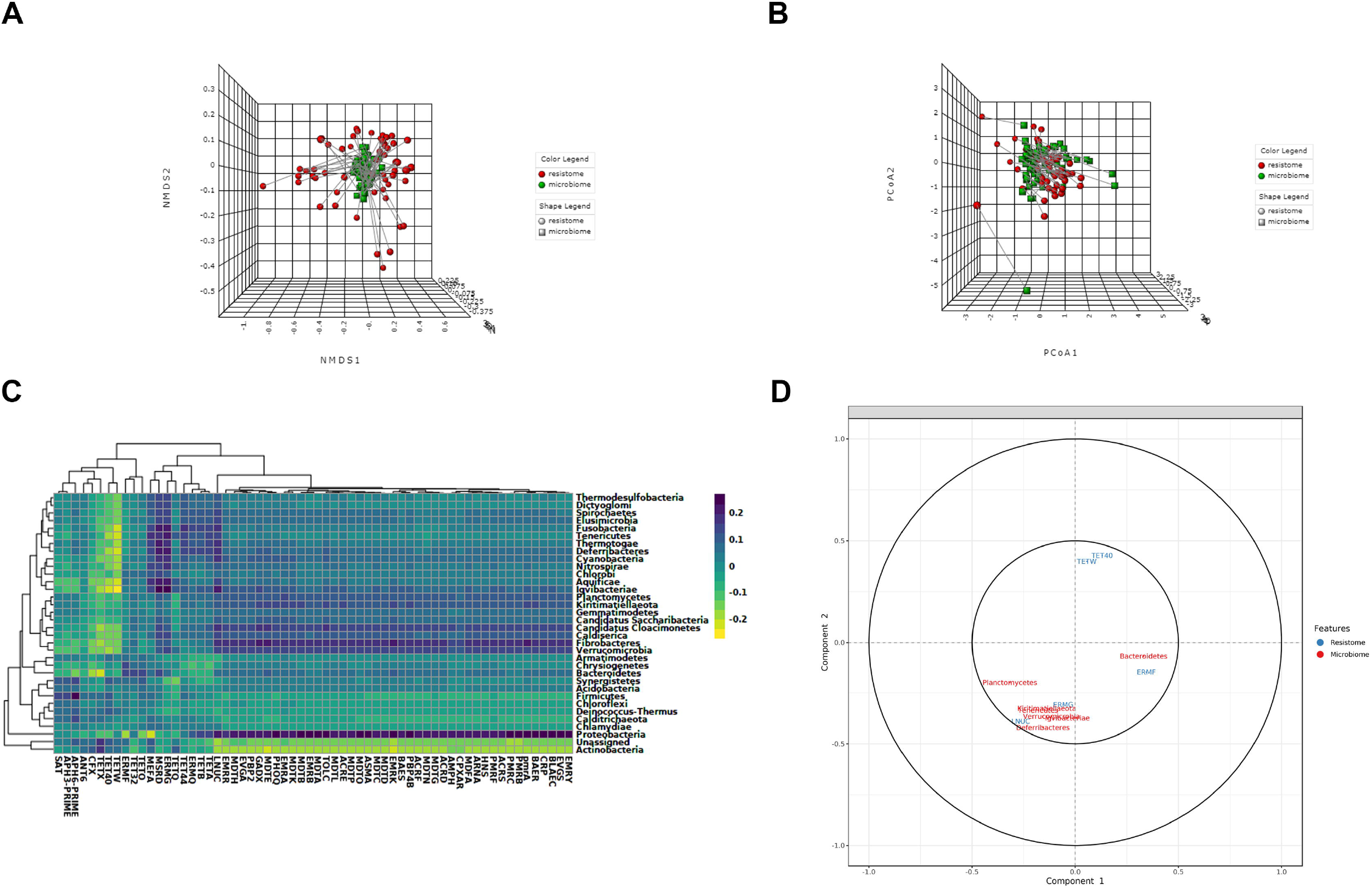
Example outputs from Integration module of ResistoXplorer. (A) A 3D NMDS plot from Procrustes analysis with samples shape and color with regards to datasets. (B) A 3D PCoA plot from Coinertia analysis, with the length of lines connecting two points indicates the similarity of samples between two datasets. (C) A clustered image heatmap showing the correlations between and among taxa (phylum level) and ARGs (group level). (D) A correlation circle plot showing the correlation structure of features (taxa/ARGs) present in two datasets.

#### Omics data integration approaches

ResistoXplorer offers other multivariate projection-based exploratory approaches such as regularized canonical correlation analysis (rCCA) and sparse partial least square (sPLS) for the integration of microbiome and resistome data. These approaches aim at highlighting the correlations between high dimensional ‘omics’ datasets. They are implemented primarily based on the R mixOmics package [64]. By default, both the datasets are normalized during such analyses using their recommended normalization technique (*i.e*., clr transformation). Users can choose or tune the number of components and regularization parameters for rCCA. In the case of sPLS, all variables are selected in both datasets by default. In addition to sample plots, other variable plots like clustered image maps (Figure 5C) and correlation circle plots (Figure 5D) are displayed to facilitate the interpretation of the complex correlation structure between datasets.

#### Pairwise microbe-ARG correlation analysis

This section enables users to determine if there are strong relationships (co-occurrence patterns) between individual microbial taxa (microbiome) and ARGs (resistome) using univariate correlation analysis. Users can perform such analysis using four different types of classical and more recent approaches, including Spearman, Pearson, CCLasso and Maximal Information Coefficient. Since the Spearman correlation analysis is most commonly used in resistome studies [65] and seems to perform overall better than other approaches in identifying pairwise associations between omics data, it has been present as a first choice in ResistoXplorer [66]. In particular, features (ARGs or taxa) that are not present in half of the samples across all the groups are removed to eliminate an artificial association bias before performing Spearman correlation analysis [65, 67]. By default, these approaches (except CCLasso) use the microbiome and resistome relative abundances (proportions) for correlation inference. While the CCLasso is based on log-ratio normalization of microbiome and resistome compositional data. Due to the lack of consensus on each approach in different conditions, it is recommended to compare results from multiple methods [66, 44]. The analysis can be conducted on taxa at their different taxonomic annotations (*i.e*., phylum, genus and species) and on resistome at different functional levels (e.g., class, mechanisms, ARG, etc.) based on available annotations. Users can select strong and significant pairwise correlations using a combination of absolute correlation coefficients and adjusted P-value. The results are represented as a co-occurrence network (Figure 6A) with each node indicating either a microbial taxon or a resistance determinant (ARG). The nodes can be sized based on their network topological measures (degree and betweenness). Users can double click on a node to highlight its corresponding correlated nodes in a network. The width and color of an edge indicate the strength and direction of the correlation between two nodes. The nodes are colored as well as shaped according to the dataset. The underlying correlation matrices and network centrality-based measures are also available to download.

**Figure 6:**
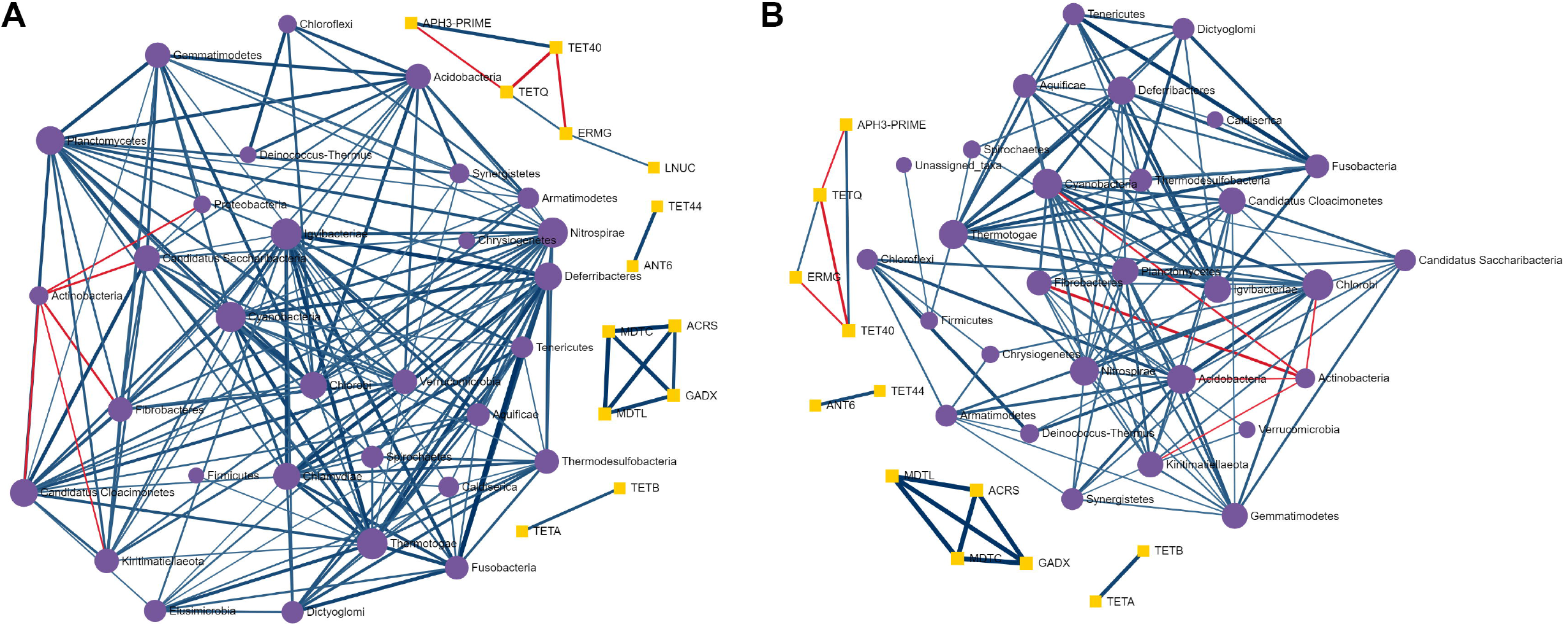
Illustration of pairwise correlation analysis results. (A) and (B) A co-occurrence network showing the strong and significant pairwise correlations between taxa (phylum level) and ARGs (class level) identified using Spearman and Pearson correlation analysis, respectively.

### ARGs-microbial host network exploration

The module offers users the possibility to understand the complex ‘multiple-to-multiple’ relations between ARGs and microbial hosts, using an advanced network-based visual analytics system. It is straightforward to identify key players from a network perspective, for instance, by looking for those ARGs that are found in multiple microbes or by identifying those microbes that simultaneously contain multiple ARGs of interest.

#### ARG-microbial host association data collection

ResistoXplorer currently supports four reference databases (ResFinder [33], CARD [32], ARDB [39] and BacMet [41]) and a recently published AMP dataset [42] for network-based microbial host associations exploration of ARGs. These primary databases contain direct or indirect information regarding the microbial host for the reference ARGs. In the latter case, the information on microbial host associated with each of the reference ARGs has been collected from their corresponding GenBank Accession number using a combination of text mining and manual curation, like in ResFinder and ARDB. Further, this information has been manually annotated to improve name readability and remove redundancy. Moreover, the available functional annotation information of ARGs was also collected and organized into sets to facilitate enrichment analysis.

#### ARG microbial host association table and network creation

The uploaded list of ARGs are searched against the selected target database. This list can comprise significant ARGs detected in differential abundance testing or ARGs identified through high-throughput qPCR. The results will be represented as an association table with each row corresponding to a particular reference ARG (sequence) and its potential microbial host. When available, the table also provides other association information along with hyperlinks to the corresponding GenBank Accession number and PubMed literature. Users can directly remove each row by clicking the delete icon in the last column to keep only high-quality associations supported by literature or experimental evidence. The resulting ARG-host associations are used to build the default networks. Since not all the nodes will be connected, this approach may lead to the generation of multiple networks. The statistics of nodes and edges are provided for users to have an overview of the size and complexity of the generated networks. Further, users can also filter the nodes based on their topological measures (degree and betweenness) in case of large networks for better interpretation.

#### Network visualization and functional analysis

The resulting networks are visualized using HTML5 canvas and JavaScript-based powerful and fully-featured visualization system. This system is implemented based on a previously published visual analytics tool [68]. It is comprised of three main components - the central network visualization area, the network customization and functional analysis panel on the left, and the right panel containing a node table (Figure 7). Users can intuitively visualize and manipulate the network in the central area using a mouse with a scroll wheel. For example, users can scroll the wheel to zoom in and out the network, hover the mouse over any node to view its name, click a node to display its details on the bottom-right corner or double click a node to select it. The horizontal toolbar to the top exhibits basic functions to manipulate the network. The first is the color picker to enable users to choose a highlight color for the next selection. Users can also select and drag multiple nodes by using the dashed square icon in the toolbar.

**Figure 7:**
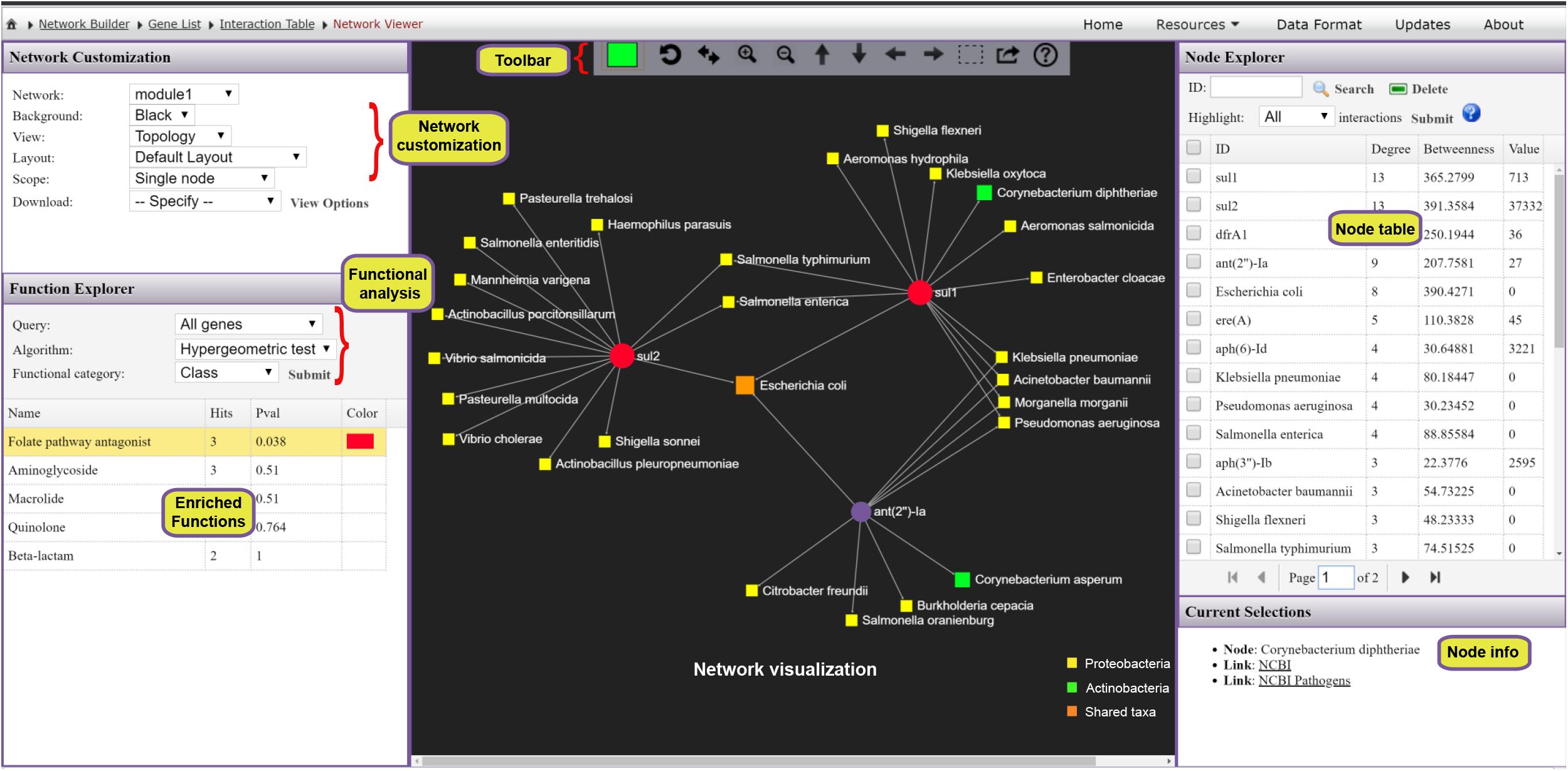
A screenshot of ResistoXplorer network visual analytics system. The view is divided into three main compartments with the network visualization (toolbar on top) at the center, the node table on the right and the network customization panel together with functional annotation table on the left. Users can easily highlight and manually organize different groups of nodes based on either their annotations or connectivity patterns. It is straightforward to identify those ARGs that are found in multiple microbes, or those microbes that simultaneously contain multiple ARGs of interest.

The network customization panel provides various options to configure the general visualization features of the default network or to specify the range of mouse operation. The ‘Layout’ option enables users to perform automatic network layout using different algorithms; the ‘Background’ option enables users to select between a white and black background. The range of mouse operations during highlighting and dragging-and-dropping can be varied using the ‘Scope’ option. For instance, in ‘single node’ mode, only the node which has been clicked or dragged will be highlighted or affected, whereas all the nodes that are being selected by the users will be affected in the ‘all highlights’ mode. Additionally, the ‘Download’ option allows users to either save the current network in different formats or to download the network file in GraphML format for visualization in other software. The node table on the right panel displays the ARGs and their microbial hosts along with corresponding network topological measures. The corresponding abundance values, if available, will also be presented in the last column. Users can directly click on any row of interest to select, and the network view will automatically zoom to the related node. The bottom right panel provides more detailed info related to the node(s) being highlighted or currently selected on the network.

ResistoXplorer also supports functional enrichment analysis of the ARGs present within the current network using hypergeometric tests. This approach coupled with the network visualization system has the potential to provide better interpretation of AMR resistance mechanisms and inform on possible dissemination routes of ARGs. The categories or levels at which enrichment analysis can be performed will be based on the initially selected database. The enrichment analysis results are shown as a table on the left panel. By clicking on a row of the result table, users can highlight all nodes related to an enriched function within the network. There are also several other options and functionalities present, which allow users to intuitively explore, manipulate and customize the ARG-microbial host association networks.

### Use Case

To illustrate the functionality of the tool, we have selected two recent resistome studies with publicly available metagenomic datasets [69, 70]. These datasets have been mounted as an example sets in the ‘ARG Table’ module of our tool. In Doster et al., 2018 [70], the authors have examined the effects of tulathromycin (antimicrobial drug) on gut microbiome and resistome using commercial feedlot cattle, which we describe here in more detail. Two groups of cattle were used, with one untreated group, and the other treated with metaphylaxis. Fecal samples were collected from 15 cattle within each group at two time points-baseline (Day 1) and after 11 days (Day 11). Shotgun sequencing was performed on the extracted metagenomic DNA, and reads were aligned to MEGARes and custom Kraken database for resistome and microbiome characterization by the authors. The resulting resistome abundance profile is uploaded to the ARG Table module of ResistoXplorer for further downstream analysis and exploration. Since the reads were annotated using the MEGARes database, we have directly selected the precompiled functional annotation information of the corresponding database to annotate and classify ARGs (gene accessions) at higher functional levels. All the ARGs are mapped and classified at three functional hierarchical levels - class, mechanism and group as per the MEGARes classification scheme. We first compared the resistome alpha diversity at the mechanism level; the Shannon diversity indices of the treatment group decrease over time, but the diversity changes are not prominent in the untreated group (Figure 2B). The composition profiling is carried out to explore and represent the gut resistome of cattle. As shown in Figure 2A, the resistome composition at the class level is dominated mainly by the ARGs that confer resistance to tetracycline and the macrolide-lincosamide-streptogramin (MLS) class of antibiotics in all the samples. The hierarchical composition profiling using the Sankey diagram showed that almost all of the ARGs that belongs to tetracycline-class confer resistance through ribosomal protection proteins. In contrast, most of the ARGs that belong to the MLS-class confer resistance through macrolide efflux pumps. Moreover, the resistome composition at different functional levels was quite similar between treated and untreated cattle (Figure 2C). The ordination analysis at AMR mechanism level using PCA (CoDA-based) and PCoA indicated that the resistome composition of treated and untreated groups were significantly different between time points (ANOSIM: R = 0.49, P-value < 0.05; Figure 3B and 3C). We also performed differential abundance testing on fecal resistome profile using metagenomeSeq and ALDEx2 (CoDA-based) at class, mechanism, and group level. No significant features were found to be differentially abundant between treatment groups using both these methods at all levels. All these analyses confirm and replicate the previous findings and results of the original publication. We also performed additional analyses of the data to highlight the utility and exploratory capabilities of ResistoXplorer. The rarefaction curve analysis indicated that enough sequencing depth was achieved to describe the ARG richness in all the metagenomic samples (Figure 2D). The heatmap showed that most of the features (except TETQ, MEFA, TETW and TETO) at group level have a very low abundance and sparse representation across all the 60 samples. The distinct abundance pattern of TET40 group is observed when comparing the Day1 with Day11 for both the treatment groups (Figure 4A). The hierarchical clustering analysis showed that samples belonging to both treated and untreated groups are clustered effectively based on time points (Figure 4B). The ARGs belonging to tetracyclines and MLS classes comprise the core resistome in cattle, based on their abundance and prevalence level (Figure 3A). Additionally, both Random Forest and SVM algorithms suggested that the treatment groups could not be predicted with high accuracies based on the resistome profiles of fecal samples, confirming the findings that tulathromycin does not seem to influence the gut resistome in cattle (Figure 4C and 4D).

Furthermore, the bacterial and ARG abundance profiles are uploaded to the Integration module of ResistoXplorer to explore the relationship between the fecal microbiome and resistome in cattle. The application of Procrustes analysis suggested that there were no significant overall similarities between the resistome and bacterial abundance profile (M2 = 0.23, P-value > 0.05; Figure 5A). However, the results from Coinertia analysis indicated that resistome and bacterial composition are moderately correlated with statistical significance (RV coefficient = 0.47, P-value < 0.05; Figure 5B). To deal with the result discrepancies between approaches, we also investigated the pairwise associations between individual taxa (microbiome) and ARGs (resistome) using Spearman and Pearson correlation analysis. The results showed that no significant and strong pairwise correlations (criteria: absolute correlation coefficient > 0.7; adjusted P-value < 0.05) were identified between any taxa (phylum level) and ARG (group level) (Figure 6A and 6B), suggesting that the gut resistome was not correlated and structured by bacterial composition.

### Comparison with Other Tools

A variety of tools or pipelines have been developed in recent years to support resistome analysis of metagenomic data [15, 33, 36, 37, 71, 72]. Most of these tools have been designed primarily for raw reads processing and annotation, with limited or no support for interactive visual exploration and downstream analysis. ResistoXplorer complements these tools and resources by providing real-time visual analytics experience along with comprehensive support for statistical, visual and exploratory analysis on the metagenomic resistome data. AMR++ Shiny [15] is a web-based R/shiny application dedicated for basic exploratory and statistical analysis of metagenomic resistome and microbiome data. More recently, resistomeAnalysis [67] R package also supports visualization, comparative and integrative analysis of resistome abundance profile. Based on the detailed comparisons among those tools (Table 1), it is clear that ResistoXplorer offers a unique set of features and functions with regards to comprehensive statistical and exploratory data analysis, visualization, integrative data analysis and ARG-microbial network visual analysis.

**Table 1:** Comparison of ResistoXplorer with other web-based tools (except resistomeAnalysis R package) supporting downstream analysis of metagenomic resistome data. Tools exclusively dedicated for sequence annotations are not included. The URL of each tool is provided under the table.

| **Tools** | **ResistoXplorer** | **AMR++ Shiny** | **resistomeAnalysis** | **WHAM!** |
| --- | --- | --- | --- | --- |
| Platform | Web-based | Web-based + locally installable | R package | Web-based |
| Registration | No | No | - | No |
| **Data Processing** |  |  |  |  |
| Data Input | Abundance tables | Abundance tables | Abundance tables | Abundance tables (Biobakery + EBI) |
| Functional annotation | User-defined + collected from >10 AMR databases | MEGARes | CARD | User-defined |
| Filtering | Abundance, variance | Abundance (Quantile) | - | Variance |
| Normalization | Scaling, transformation, rarefying | Cumulative Sum Scaling (CSS), rarefying | Total Sum Scaling (TSS), proportion | Proportion, clr |
| **Composition Profiling** |  |  |  |  |
| Visual profiling | Stacked bar plot, stacked area, sankey diagram, zoomable sunburst, treemap | Stacked bar plot | Stacked bar plot | Interactive Stacked bar plot |
| Alpha diversity | Multiple | - | Richness | - |
| Ordination analysis | PCoA, NMDS & PCA (2D & 3D) | PCA & NMDS (Bray-Curtis) (2D) | PCoA (2D) | - |
| **Comparative analysis** |  |  |  |  |
| Differential analysis | DESeq2, metagenomeSeq, EdgeR, LefSe, ALDEx2, ANCOM | metagenomeSeq | DESeq2 | ALDEx2 |
| Classification | Random Forests, SVM | - | - | - |
| Other functions | Heatmaps, dendrogram, correlation, core resistome, alpha rarefaction curves | Heatmaps, Alpha rarefaction bar plots | Heatmaps, dendrogram, correlation, core resistome | Interactive Heatmaps, correlation |
| **Integrative analysis** | Procrustes, Coinertia,<br>rCCA, sPLS,<br>Spearman, Pearson,<br>CCLasso, MIC | - | Spearman | Visual<br>comparisons |
| **Gene (ARG) List**<br>**exploration** | ARG-microbial host<br>network visual<br>analytics & functional<br>analysis | - | - | - |

ResistoXplorer: http://www.resistoxplorer.no
AMR ++ Shiny: https://github.com/lakinsm/amrplusplus-shiny
resistomeAnalysis (R package): https://github.com/blue-moon22/resistomeAnalysis
WHAM!: https://ruggleslab.shinyapps.io/wham_v1/

### Limitations and Future Directions

The ARG table module can be used for visualization and analysis of resistome profiles characterizing different genetic determinants present within an AMR reference database. However, ResistoXplorer does not allow users to choose multiple precompiled functional annotation databases to analyze the entire ecologies (antimicrobial drugs, biocide, metal and other resistance drivers) of resistance determinants. More importantly, the functional annotations collected in ResistoXplorer mainly depend on the information and classification scheme present in the original databases. Hence, there might still be some acyclic and hierarchical functional annotation structure in databases, which users need to curate for accurate count-based analyses. The biocuration of ARGs and their functional annotation structure for the supported databases is beyond the scope of the study. Although some of the supported databases are no longer updated, such as ARDB and ARG-ANNOT [5, 6], excluding them would have limited the possibility of exploring and analyzing previous datasets, as well as a variety of present studies that still use them. In regard to the ARG-microbial host associations module, these are limited by the type and quality of information available in the databases. Currently, ResistoXplorer offers limited functionalities and features for vertical data integration and pairwise correlation analysis. As most of the advanced approaches for performing this type of analysis on multidimensional datasets are based on computationally intensive re-sampling (cross-validation) and permutation-based approaches to calculate statistical significance, which adds layers of complexity and computational power demands, that is often challenging for a real-time interactive web application. In future versions, we plan to continuously update and expand our database and analysis support for exploration of mobilome, virulome and other resistance determinants of relevance in AMR metagenomic studies.

## CONCLUSION

Whole metagenomic sequencing studies are providing unparalleled knowledge on the diversity of resistomes in the environment, animals and humans, and on the impact of interventions, such as antibiotic use [7–12, 67, 69]. Currently, such studies and data analyses are mainly exploratory in nature. In spite of the continuous development of many new statistical approaches, there is no exclusive method that unfailingly performs well, as demonstrated by several benchmarking studies [20, 25, 26, 46]. Indeed, it has been recently suggested that metagenomic analysis should be explored comparatively using different available approaches [31]. However, this is a time consuming task and requires knowledge and bioinformatics training on the implementation of each statistical method employed. Therefore, it is critical to assist researchers and clinical scientists in the field to easily explore their own datasets using a variety of approaches, in realtime and through interactive visualization, to facilitate data understanding and hypothesis generation. ResistoXplorer meets these requirements by offering comprehensive support for composition profiling, statistical analysis, integrative analysis and visual exploration of resistome data. Conversely, such analysis is entirely dependent on the comprehensiveness and quality of the AMR reference databases [5, 6]. Hence, the use of continuously updated and curated databases with a simple, acyclic and hierarchical functional annotation scheme is desired for accurate downstream analysis. Lastly, ResistoXplorer will continuously be updated to follow the advancements in approaches for resistome analysis. We believe ResistoXplorer will have the potential to find large applicability as a useful resource for researchers in the field of AMR.

## ACKNOWLEDGEMENTS

This work has been supported and used the computing resources at the University of Oslo.

